# Starvation-induced regulation of carbohydrate transport at the blood-brain barrier is TGF-β-signaling dependent

**DOI:** 10.1101/2020.09.21.306308

**Authors:** Helen Hertenstein, Ellen McMullen, Astrid Weiler, Anne Volkenhoff, Holger M. Becker, Stefanie Schirmeier

## Abstract

During hunger or malnutrition animals prioritize alimentation of the brain over other organs to ensure its function and thus their survival. This so-called brain sparing is described from Drosophila to humans. However, little is known about the molecular mechanisms adapting carbohydrate transport. Here, we used Drosophila genetics to unravel the mechanisms operating at the blood-brain barrier (BBB) under nutrient restriction. During starvation, expression of the carbohydrate transporter Tret1-1 is increased to provide more efficient carbohydrate uptake. Two mechanisms are responsible for this increase. Similarly to the regulation of mammalian GLUT4, Rab-dependent intracellular shuttling is needed for Tret1-1 integration into the plasma membrane, even though Tret1-1 regulation is independent of insulin signaling. In addition, starvation induces transcriptional upregulation controlled by TGF-β signaling. Considering TGF-β-dependent regulation of the glucose transporter GLUT1 in murine chondrocytes, our study reveals an evolutionarily conserved regulatory paradigm adapting the expression of sugar transporters at the BBB.

## Introduction

A functional nervous system is essential for an animal’s survival. To properly function, the nervous system needs a disproportionately large amount of energy relative to its size. The human brain for example accounts for only about 2% of the body’s weight but uses around 20% of the resting oxygen consumption (Laughlin et al., 1998). Similarly, the insect retina consumes approximately 10% of the total ATP generated (Harris et al., 2012; Laughlin et al., 1998; Mink et al., 1981).

The nervous system is very susceptible to changing extracellular solute concentrations and thus needs to be separated from circulation. This task is performed by the blood-brain barrier (BBB), which prevents paracellular diffusion, and thereby uncontrolled influx of ions, metabolites, xenobiotics, pathogens and other blood-derived potentially harmful substances. Protein, ion and metabolite concentrations fluctuate much stronger in circulation than in the cerebrospinal fluid, the brains extracellular milieu (Begley, 2006). Thus, fluxes over the BBB must be tightly regulated and only small lipid soluble molecules and gases like O_2_ and CO_2_ can diffuse freely (van de Waterbeemd et al., 1998).

The enormous energy demand of the nervous system is mainly met by carbohydrate metabolism. The human brain takes up approximately 90 g glucose per day during adulthood, and up to 150 g per day during development (Kuzawa et al., 2014). Since glucose and other carbohydrates are hydrophilic molecules, free diffusion over the BBB is impossible. Therefore, carbohydrates need to be transported into the nervous system via specialized transport proteins. In mammals, Glucose transporter 1 (GLUT1, encoded by the *Slc2a1* (*solute carrier family 2 member 1*) gene) is considered to be the main carbohydrate transporter in the BBB-forming endothelial cells. Aberrations in carbohydrate availability or transport are thought to be a major factor in the development of diverse neurological diseases such as Glut1 deficiency syndrome, Alzheimer’s disease or epilepsy (Arsov et al., 2012; Hoffmann et al., 2013; Kapogiannis and Mattson, 2011; Koepsell, 2020). Therefore, understanding the regulatory mechanisms that govern carbohydrate transport into the nervous system is of major interest. Interestingly, it has been reported that endothelial GLUT1 expression is increased upon hypoglycemia (Boado and Pardridge, 1993; Kumagai et al., 1995; Simpson et al., 1999, reviewed in Patching, 2016; Rehni and Dave, 2018). However, the molecular mechanisms that control this upregulation are not yet understood. In addition, upon oxygen or glucose deprivation that are a consequence of ischemia, expression of the sodium glucose cotransporters SGLT1 (*Slc5a1*) and SGLT2 (*Slc5a2*) is induced in brain endothelial cells (Elfeber et al., 2004; Enerson and Drewes, 2006; Nishizaki et al., 1995; Nishizaki and Matsuoka, 1998; Vemula et al., 2009; Yu et al., 2013). Overall, this indicates that carbohydrate transport at the BBB can adapt to changes in carbohydrate availability in various ways. However, the molecular underpinnings of the different regulatory processes are still elusive.

Similarly to vertebrates, the insect nervous system must be protected by a BBB. Since insects have an open circulatory system the brain is not vascularized but is surrounded by the blood-like hemolymph. In Drosophila the BBB surrounds the entire nervous system to prevent uncontrolled entry of hemolymph-derived substances. It is formed by two glial cell layers, the outer perineurial and inner subperineurial glial cells (reviewed in Limmer et al., 2014; Yildirim et al., 2019). The Drosophila BBB shares fundamental functional aspects with the vertebrate BBB. The subperineurial glial cells build a diffusion barrier by forming intercellular pleated septate junctions that prevent paracellular diffusion (Stork et al., 2008). In addition, efflux transporters export xenobiotics and many solute carrier (SLC) family transporters supply the brain with essential ions and nutrients (Desalvo et al., 2014; Hindle and Bainton, 2014; Lane and Treherne, 1972; Mayer and Belsham, 2009; Stork et al., 2008; reviewed in Weiler et al., 2017). In the Drosophila hemolymph, in addition to glucose, trehalose, a non-reducing disaccharide consisting of two glucose subunits linked by an α,α-1,1-glycosidic bond, is found in high quantities. Fructose is also present albeit in low and highly fluctuating concentrations, making its nutritional role questionable (Blatt and Roces, 2001; Broughton et al., 2008; Lee and Park, 2004; Pasco and Léopold, 2012; Wyatt and Kalf, 1957). Transcriptome data of the BBB-forming glial cells suggests expression of several putative carbohydrate transporters (Desalvo et al., 2014; Ho et al., 2019). The closest homologues of mammalian GLUT1-4 are dmGlut1, dmSut1, dmSut2, dmSut3 and CG7882. dmGlut1 has been shown to be expressed exclusively in neurons (Volkenhoff et al., 2018). *In situ*, microarray and single cell sequencing data indicate very low or no expression for dmSut1-3 and CG7882 in the nervous system (Croset et al., 2018; Davie et al., 2018; Weiszmann et al., 2009). The carbohydrate transporter Tret1-1 (Trehalose transporter 1-1) is specifically expressed in perineurial glia (Volkenhoff et al., 2015). Tret1-1 is most homologous to mammalian GLUT6 and GLUT8 and has been shown to transport trehalose when heterologously expressed in *Xenopus laevis* oocytes (Kanamori et al., 2010).

The Drosophila nervous system, as the mammalian nervous system, is protected from the effects of malnutrition through a process called brain sparing. It has been shown that upon nutrient restriction neuroblasts (neural stem cells) can still divide and are thus protected from the growth defects that are caused by a lack of proper nutrition in other tissues (reviewed in Lanet and Maurange, 2014). This protection is achieved by Jelly belly (Jeb)/Anaplastic lymphoma kinase (ALK) signaling that constitutes an alternative growth promoting pathway active in neuroblasts (Cheng et al., 2011). However, if the brain continues developing and keeps its normal function, nutrient provision needs to be adapted to ensure sufficient uptake, even under challenging circumstances, like low circulating carbohydrate levels. How nutrient transport at the BBB is adapted to meet the needs of the nervous system even under nutrient restriction has not been studied.

Here, we show that carbohydrate transporter expression in Drosophila as in mammals adapts to changes in carbohydrate availability in circulation. Tret1-1 expression in perineurial glia of Drosophila larvae is strongly upregulated upon starvation. This upregulation is triggered by starvation-induced hypoglycemia as a mechanism protecting the nervous system from the effects of nutrient restriction. *Ex vivo* glucose uptake measurements using a genetically encoded FRET-based glucose sensor show that the upregulation of carbohydrate transporter expression leads to an increase in carbohydrate uptake efficiency. The compensatory upregulation of Tret1-1 transcription is independent of insulin/adipokinetic hormone signaling, but instead depends on TGF-β signaling. This regulatory mechanism that allows sparing the brain from the effects of malnutrition is likely conserved in mammals, since mammalian Glut1 is also upregulated in the BBB upon hypoglycemia and has been shown to be induced by TGF-β signaling in other tissues (Boado and Pardridge, 1993; Kumagai et al., 1995; Simpson et al., 1999; Lee et al., 2018).

## Results

### Tret1-1 is upregulated in perineurial glial cells upon starvation

The Drosophila larval brain is separated from circulation by the blood-brain barrier to avoid uncontrolled leakage of hemolymph-derived potentially harmful substances. At the same time, the blood-brain barrier also cuts off the brain from nutrients available in the hemolymph. Thus, transport systems are necessary to ensure a constant supply of nutrients, including carbohydrates. The trehalose transporter Tret1-1 is expressed in the perineurial glial cells of the larval and adult nervous system (Volkenhoff et al., 2015). In order to better understand whether carbohydrate transport at the BBB is adapted to the metabolic state of the animal, we analyzed Tret1-1 dynamics under different physiological conditions. In fed animals Tret1-1 can be found at the plasma membrane of the perineurial glial cells, but a large portion localizes to intracellular vesicles (Figure 1A, Volkenhoff et al., 2015). We subjected wild type larvae to chronic starvation applying a well-established paradigm that allows 48 h of starvation without disturbing development (Zinke et al., 2002). Starvation increases Tret1-1 protein levels in the perineurial glial cells (Figure 1A). Furthermore, an enrichment of Tret1-1 protein at the plasma membrane was observed (Figure 1A, asterisk), showing that starvation induces changes in Tret1-1 levels as well as localization.

**Figure 1.**
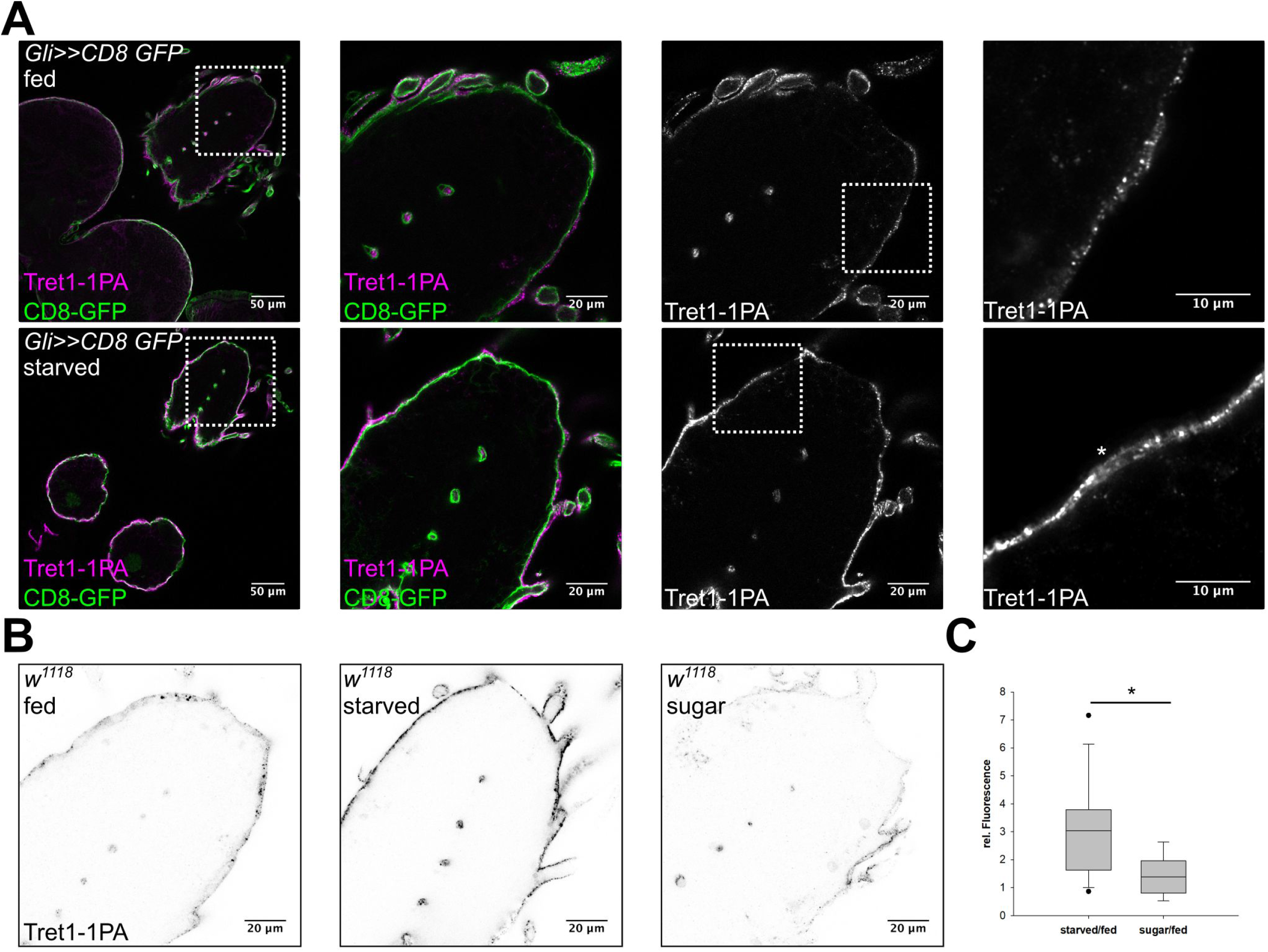

### Intracellular trafficking of Tret1-1 is Rab7 and Rab10 dependent

Three mammalian Glucose transporters, GLUT4, GLUT6 and GLUT8, are regulated via trafficking between storage vesicles and the plasma membrane (Corvera et al., 1994; Cushman and Wardzala, 1980; Lisinski et al., 2001; Suzuki and Kono, 1980). Similarly, a large amount of Tret1-1 localizes to intracellular vesicles (Figure 1A). Thus, intracellular trafficking of Tret1-1 may partially regulate carbohydrate uptake into the perineurial glial cells.

To analyze if regulation of Tret1-1 expression requires intracellular trafficking, we studied the involvement of different Rab-GTPases. Utilizing an EYFP-Rab library available for Drosophila (Dunst et al., 2015) we found that subsets of Tret1-1 positive vesicles are also positive for Rab7, Rab10, Rab19 and Rab23 (Figure S1). Rab7 is needed for the formation of late endosomes and their fusion with lysosomes, while Rab10 has been implicated in GLUT4 storage vesicle trafficking in mammals (reviewed in Guerra and Bucci, 2016; Huotari and Helenius, 2011; Klip et al., 2019). The roles of Rab19 and Rab23 are less well understood. Rab23 has been implicated in planar cell polarity and in Hedgehog regulation in response to dietary changes, but its exact functions are unclear (Çiçek et al., 2016; Pataki et al., 2010). Rab19 has been described to act in enteroendocrine cell differentiation, but its role in this process is unknown (Nagy et al., 2017).

To determine a possible functional role of these Rab-GTPases in regulating Tret1-1 trafficking, we analyzed Tret1-1 localization in the background of a glia-specific knockdown (or expression of dominant-negative forms) of the respective Rab proteins (Figure 2A,B). Silencing of Rab19 or Rab23 did not induce any misregulation or mislocalization of Tret1-1 in perineurial glial cells (data not shown). In contrast, interfering with Rab7 or Rab10 function induced distinct abnormal phenotypes (Figure 2). Panglial and BBB-glia cell-specific knockdown of Rab7 using RNA interference or expression of a dominant-negative form of Rab7, Rab7^T22N^, reduced the levels of Tret1-1 (Figure 2). The dominant-negative Rab-constructs used here are tagged with an N-terminal YFP and thus induce a weak background staining in all glial cells (Figure 2B, asterisks). The reduced Tret1-1 level in Rab7 loss of function indicates that blocking late endosome to lysosome maturation and thus possibly blocking Tret1-1 degradation, induces a negative feedback that reduces Tret1-1 expression.

**Figure 2.**
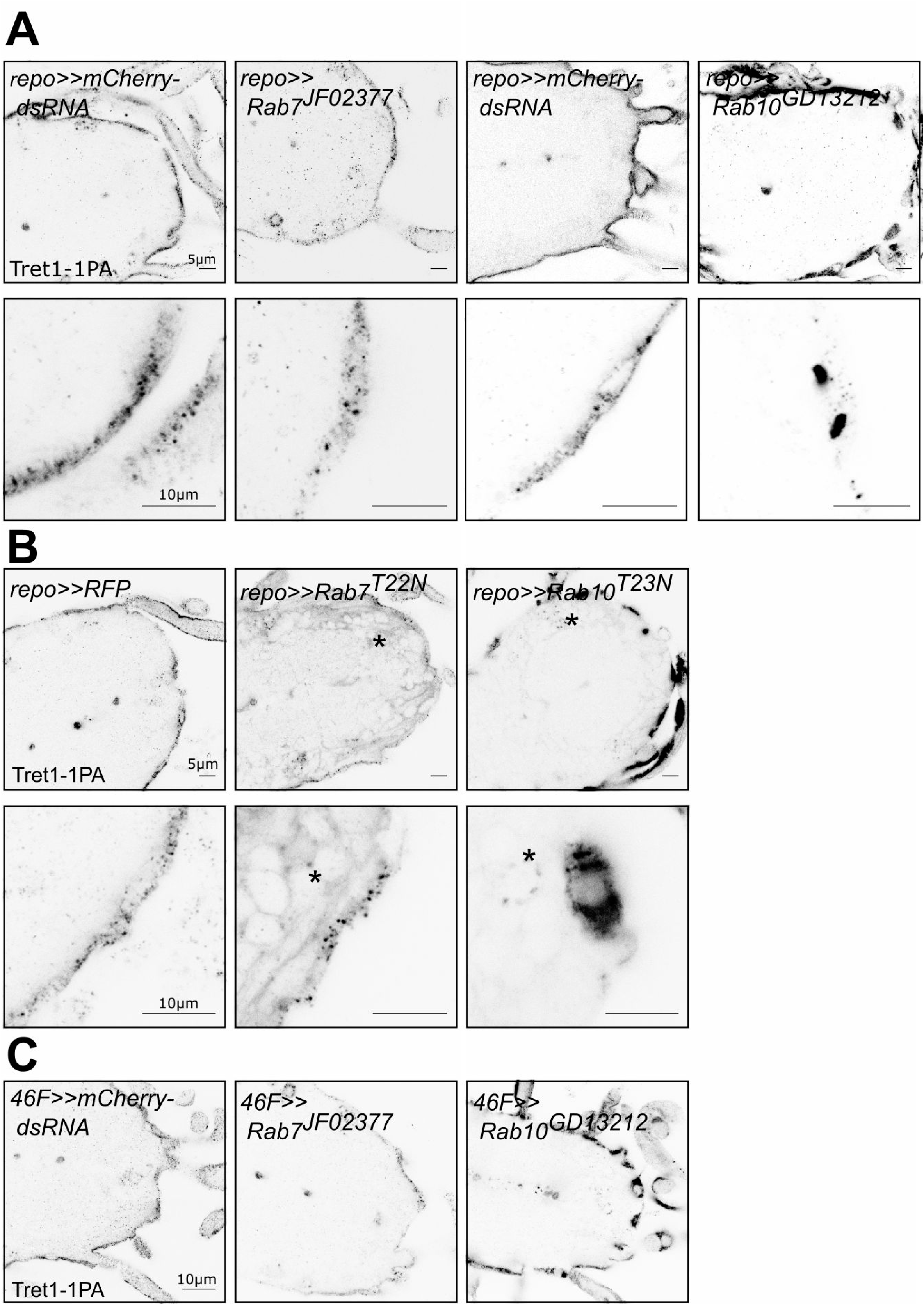

In contrast to Rab7, knockdown of Rab10 in all glia, or in the BBB-glial cells specifically, leads to a prominent accumulation of Tret1-1 in the perineurial cytosol (Figure 2). This phenotype was reproduced when a dominant-negative form of Rab10, Rab10^T23N^, was expressed in glial cells, suggesting a major role of Rab10 in delivering Tret1-1 to the plasma membrane of perineurial glial cells. In summary, Tret1-1 homeostasis is dependent on Rab-GTPase-mediated intracellular trafficking.

### Increase in Tret1-1 expression upon starvation is sugar-dependent

The expression of mammalian Glut1 in brain endothelial cells increases upon chronic hypoglycemia (Boado and Pardridge, 1993; Kumagai et al., 1995; Rehni and Dave, 2018; Simpson et al., 1999). In Drosophila, starvation results in hypoglycemia (Dus et al., 2011; Matsuda et al., 2015). Thus, we wondered if the increase in Tret1-1 protein levels described here might be induced by a reduction in circulating carbohydrate levels. To understand if dietary carbohydrates are sufficient to circumvent Tret1-1 induction, we compared animals fed on standard food, starved animals, and animals fed on 10 % sucrose in phosphate-buffered saline. Larvae kept on sugar-only food display comparable Tret1-1 levels as larvae kept on standard food (Figure 1B,C). Hence, dietary sugar abolishes Tret1-1 induction, indicating that other nutrients, like amino acids are not important for this signaling pathway (Figure 1B,C). Attempts to analyze Tret1-1 levels in larvae fed on a protein-only diet to study the influence of dietary amino acids were unsuccessful as larvae do not eat protein-only diet (no uptake of colored protein-only food into the intestine over 48 h, data not shown). This data suggests that Tret1-1 is upregulated in the perineurial glial cells upon starvation-induced hypoglycemia. Such an increase in Tret1-1 protein levels could be due to transcriptional regulation or posttranscriptional mechanisms interfering with translation or protein stability.

### Tret1-1 *is transcriptionally regulated upon starvation*

To test whether transcriptional regulation accounts for the strong increase in Tret1-1 protein upon starvation, we cloned the *tret1-1* promotor and established transgenic animals expressing either Gal4 or a nuclear GFP (stinger-GFP, stgGFP) under its control (Figure S2). We validated the expression induced by the promotor fragment by co-staining RFP expressed under *tret1-1-Gal4* control with the Tret1-1 antibody we generated previously (Volkenhoff et al., 2015). *tret1-1* promotor expression and Tret1-1 protein colocalize well in the nervous system (Figure S2B). We previously showed that Tret1-1 localizes to perineurial glial cells and some unidentified neurons (Volkenhoff et al., 2015). To further verify perineurial glial expression, we stained *tret1-1-stgGFP* animals for a nuclear perineurial glial marker, Apontic (Figure S2C, Zülbahar et al., 2018). Apontic and stgGFP colocalize in perineurial nuclei.

To analyze changes in *tret1-1* transcription levels, we subjected animals expressing stgGFP under the control of the *tret1-1* promotor to our starvation paradigm. Starvation induces a robust increase of stgGFP in the brains of starved larvae as quantified by Western Blot (Figure 3). These experiments show that the *tret1-1* promotor is induced upon starvation and thus Tret1-1 levels are transcriptionally adapted to the animal’s metabolic state.

**Figure 3.**
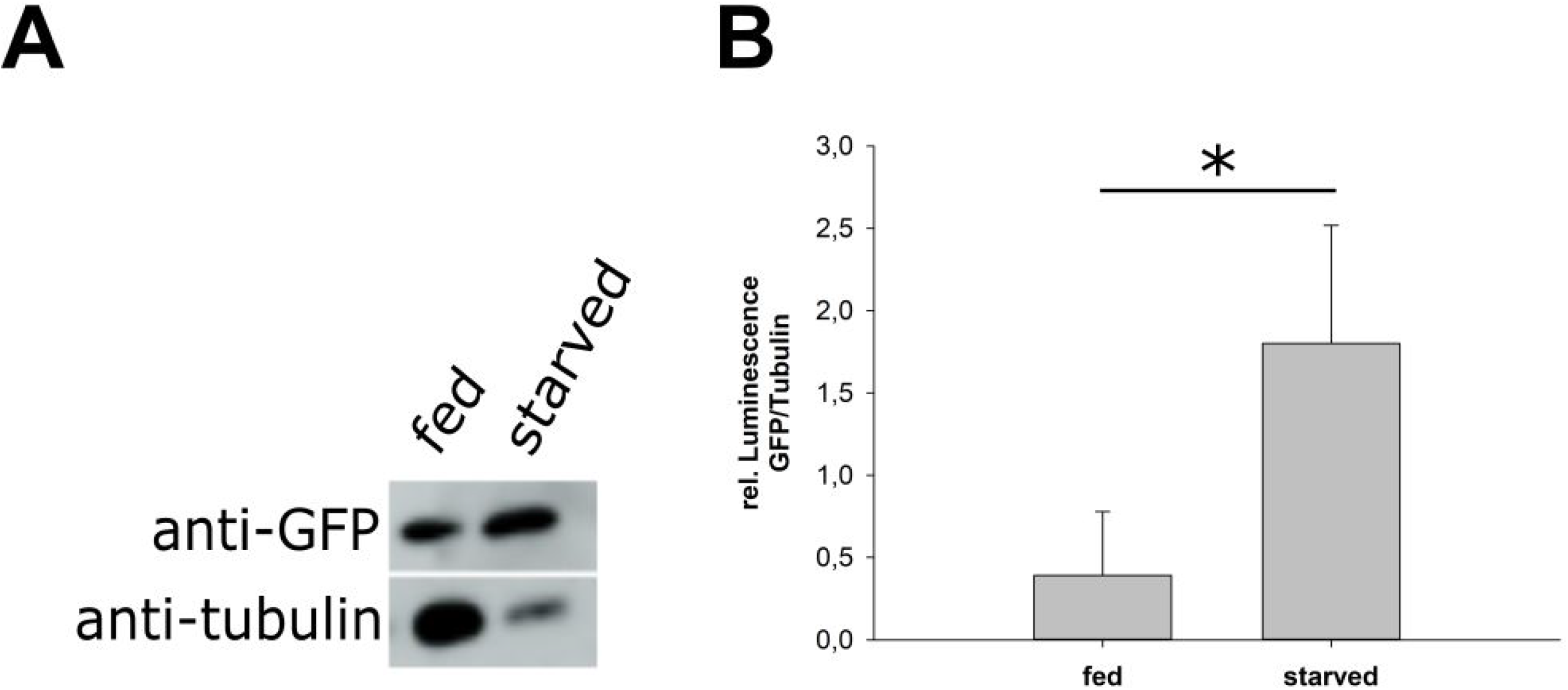

### Glucose uptake rate increases upon starvation

Tret1-1 upregulation in perineurial glial cells is most likely a mechanism that ensures efficient carbohydrate uptake into the nervous system even under conditions of low circulating carbohydrate levels. Therefore, we aimed to study the impact of Tret1-1 upregulation on carbohydrate uptake at the BBB. Kanamori et al., 2010 showed that Tret1-1 transports trehalose when heterologously expressed in *Xenopus laevis* oocytes. Since not only trehalose but also glucose and fructose are found in the Drosophila hemolymph, we analyzed whether Tret1-1 also transports other carbohydrates. Therefore, we expressed Tret1-1 in *Xenopus leavis* oocytes to study its substrate specificity. The Tret1-1 antibody is specific to the Tret1-1PA isoform, and thus at least this isoform is upregulated in the perineurial glial cells upon starvation. Therefore, we expressed a 3xHA-tagged version of Tret1-1PA in *Xenopus laevis* oocytes. The functionality of this construct was verified by its ability to rescue the lethality associated with *tret1-1^-/-^* mutants when ubiquitously expressed (using *da-Gal4*, Volkenhoff et al., 2015). Incubating *Xenopus laevis* oocytes expressing Tret1-1PA-3xHA with different concentrations of ^14^C_6_-fructose, ^14^C_6_-glucose or ^14^C_12_-trehalose for 60 min, we were able to verify the trehalose transport capacity reported previously (Kanamori et al., 2010) (Figure 4A). In addition, Tret1-1PA can facilitate uptake of glucose, while fructose is not taken up efficiently (Figure 4A).

**Figure 4.**
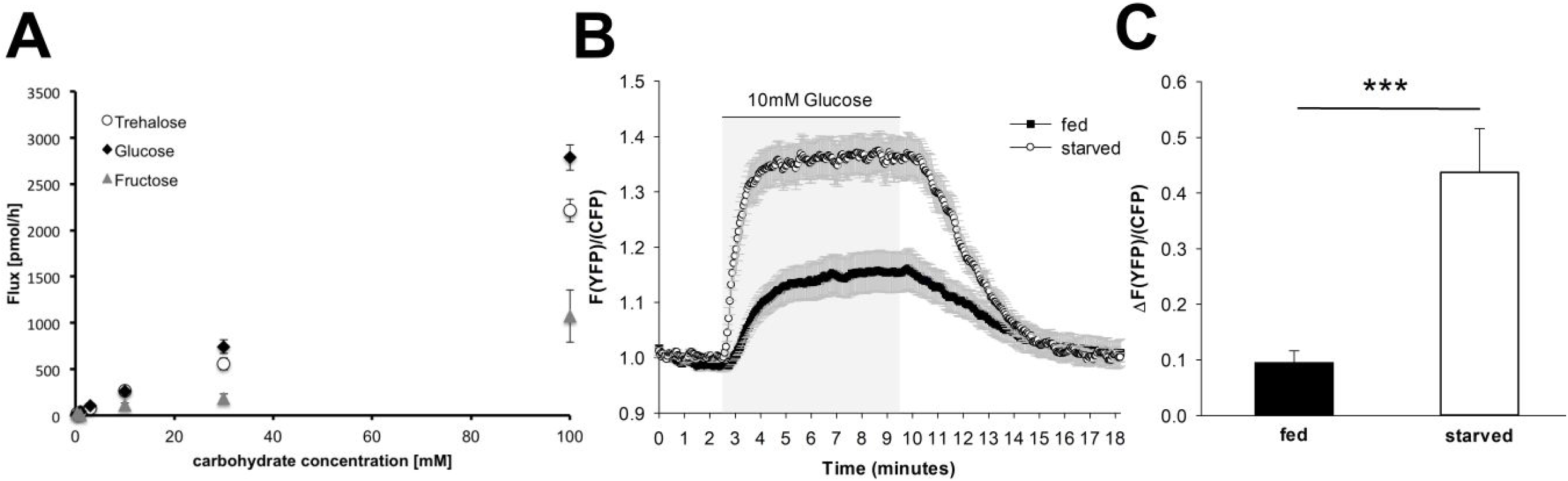

Taking advantage of the glucose transport capacity of Tret1-1, we employed the Förster resonance energy transfer (FRET)-based glucose sensor FLII^12^Pglu-700μδ6 (Takanaga et al., 2008; Volkenhoff et al., 2018) to determine the effect of Tret1-1 upregulation on carbohydrate import into the living brain. A trehalose sensor to measure trehalose uptake is unfortunately not available. However, the glucose sensor allows live imaging of glucose uptake in a cell type of choice in *ex vivo* brain preparations (Volkenhoff et al., 2018). We expressed FLII^12^Pglu-700μδ6 specifically in the BBB glial cells (9137-Gal4, Desalvo et al., 2014). The respective larvae were subjected to the starvation protocol and, subsequently, glucose uptake was measured (Figure 4B,C). The rate of glucose uptake was significantly increased in brains of starved animals compared to the brains of age-matched animals kept on standard food (Figure 4B,C). These findings show that, indeed, carbohydrate uptake into the brain is more efficient in starved animals. Such improved carbohydrate uptake most likely protects the brain from the effects of low circulating carbohydrate levels.

### Starvation-induced upregulation of Tret1-1 is insulin- and adipokinetic hormone-independent

The plasma membrane localization of mammalian GLUT4 is regulated by insulin (reviewed in Klip et al., 2019). Since starvation changes circulating carbohydrate levels, it has strong effects on insulin and adipokinetic hormone (AKH) signaling (reviewed in Nässel et al., 2015). Thus, insulin/AKH signaling may control Tret1-1 induction upon starvation. To study the implication of insulin signaling we expressed dominant-negative forms of the insulin receptor (InR, InR^K1409A^ and InR^R418P^) in the BBB-forming glial cells (Figure 5A,B). If Insulin signaling was to directly regulate Tret1-1 transcription, one would assume a negative effect, since *tret1-1* is upregulated upon starvation when insulin levels are low. If insulin signaling indeed has a negative effect on *tret1-1* expression, higher Tret1-1 levels would be expected under fed conditions upon expression of a dominant-negative InR. Expression of dominant-negative forms of InR did not changed Tret1-1 levels in fed animals in comparison to the control (Figure 5A). In addition, Tret1-1 upregulation upon starvation was indistinguishable from that observed in control animals (Figure 5A,B), indicating that Tret1-1 transcription is independent of insulin signaling.

**Figure 5.**
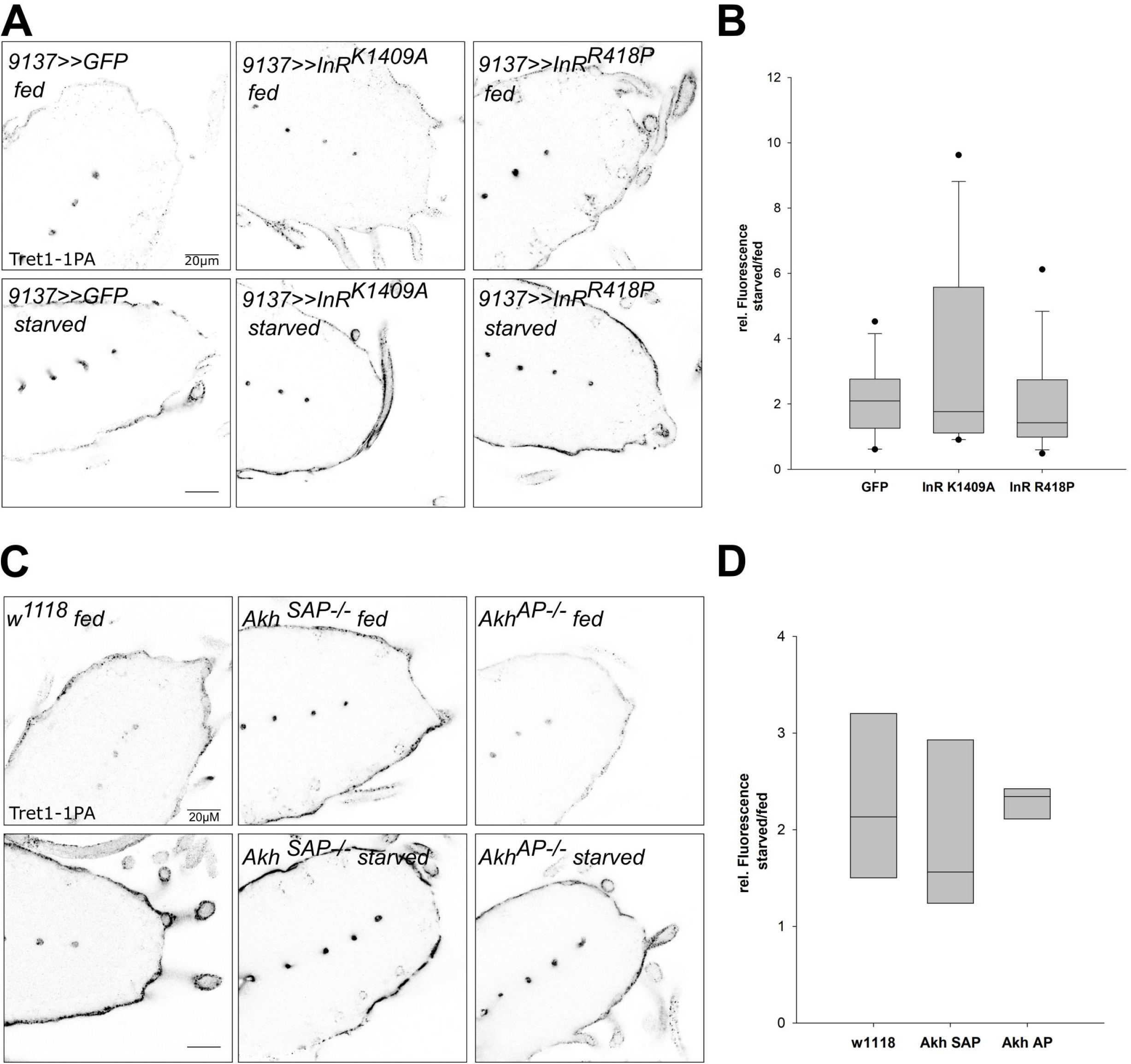

In Drosophila, AKH is thought to play a role equivalent to glucagon/glucocorticoid signaling in mammals (Gáliková et al., 2015). AKH signaling induces lipid mobilization and foraging behavior, at least in the adult animal (Gáliková et al., 2015). Thus, AKH signaling would be a good candidate to induce *Tret1-1* upregulation upon starvation. We analyzed Tret1-1 levels in *Akh^-/-^* (*Akh^SAP^* and *Akh^AP^*) mutant animals under normal conditions and starvation. Tret1-1 levels in the perineurial glial cells in both fed *Akh^-/-^* mutant larvae are indistinguishable from control levels (Figure 5C). Interestingly, Tret1-1 is still induced upon starvation in *Akh^-/-^* mutant animals (Figure 5C,D). This suggests that AKH does not play a role in Tret1-1 regulation upon starvation. In summary, the core signaling pathways regulating organismal nutrient homeostasis, Insulin and AKH signaling, are not involved in Tret1-1 upregulation upon starvation.

### Jelly belly/Anaplastic lymphoma kinase signaling does not regulate Tret1-1 expression

Tret1-1 upregulation upon starvation is likely a mechanism to spare the nervous system from the effects of restricted nutrient availability. Jelly belly (Jeb)/Anaplastic lymphoma kinase (ALK) signaling is important to allow continued developmental brain growth even upon poor nutrition (Cheng et al., 2011). To analyze if this pathway might also play a role in adapting carbohydrate transport, we knocked down *jeb* and *Alk* in all glial cells and analyzed Tret1-1 expression. *Alk* knockdown in the glial cells did not induce a Tret1-1 expression phenotype (Figure S3). Tret1-1 is still upregulated upon starvation, indicating that ALK signaling in glial cells is not involved in Tret1-1 regulation (Figure S3). *jeb* knockdown in all glial cells induced strong starvation susceptibility of the animals in our hands. Most animals died within the 48 h starvation period and analyzing Tret1-1 expression in the perineurial glial cells of escapers did not give coherent results. Nevertheless, since *Alk* knockdown shows wild typic Tret1-1 upregulation, Jeb/ALK signaling is most likely not implicated in the regulation of carbohydrate transport upon starvation.

### Transforming growth factor β signaling regulates Tret1-1 expression

In Drosophila, both TGF-β/Activin signaling and TGF-β/bone morphogenetic protein (BMP) signaling have been implicated in metabolic regulation (Ballard et al., 2010; Ghosh and O’Connor, 2014). The Activin and BMP branches of TGF-β signaling share some components, like the type II receptors Punt (Put) and Wishful thinking (Wit) and the co-Smad Medea, while other components are specific to one or the other branch (reviewed in Upadhyay et al., 2017), Figure 6C).

**Figure 6.**
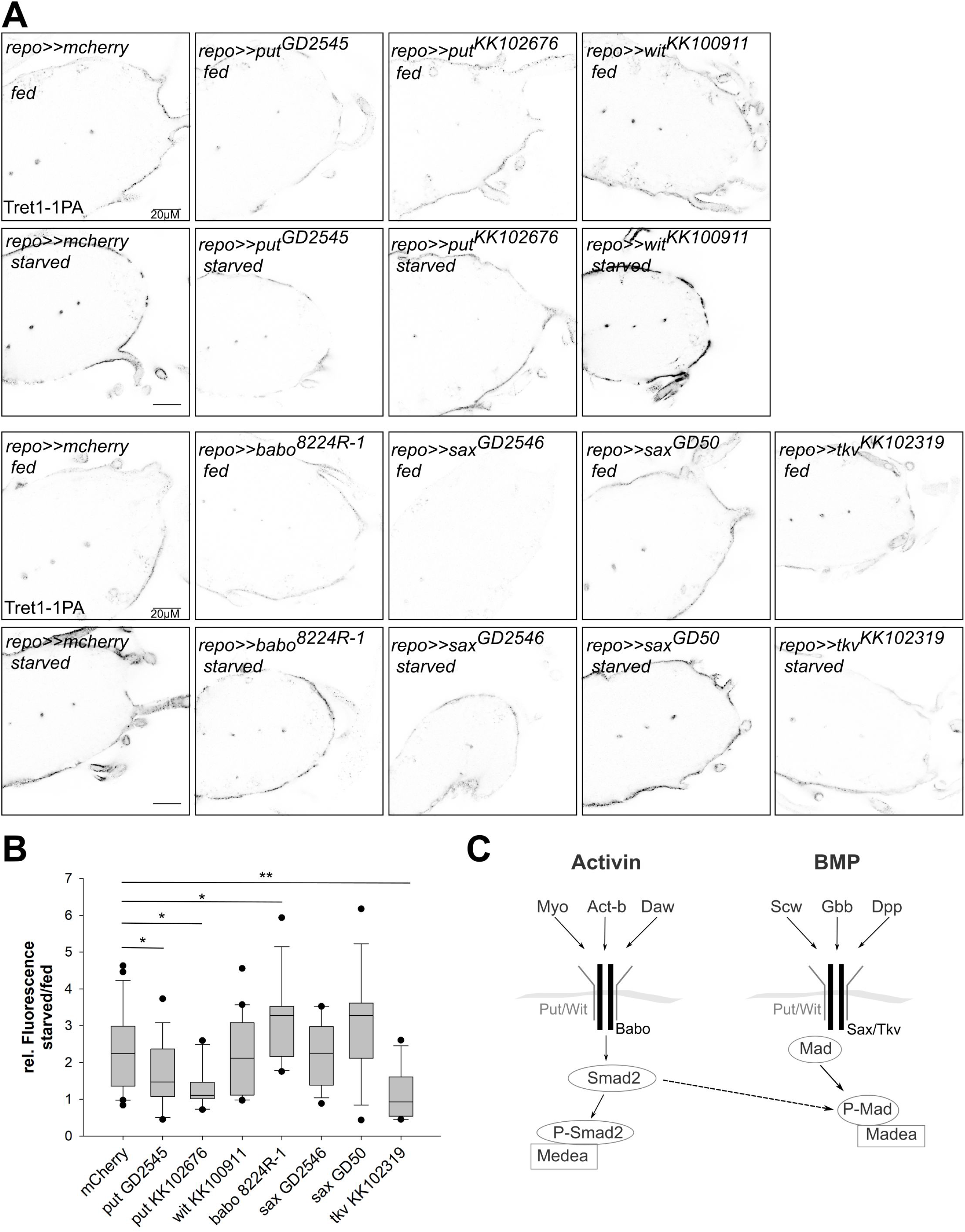

Since Put has been implicated in regulating carbohydrate homeostasis, we asked if Put-dependent TGF-β signaling could also play a role in carbohydrate-dependent Tret1-1 regulation. Thus, we expressed dsRNA constructs against *put* in a glia-specific manner and analyzed Tret1-1 levels in the perineurial glial cells of fed and starved animals (Figure 6). Indeed, starvation-dependent upregulation of Tret1-1 was completely abolished upon *put* knockdown in the glial cells using either *put^KK102676^* or *put^GD2545^*. Quantification shows no upregulation of Tret1-1 upon starvation in *put* knockdown animals (Figure 6B). In contrast, knockdown of *wit* using *wit^KK100911^*, did not affect Tret1-1 upregulation upon starvation (Figure 6). This data suggests, that Put-dependent TGF-β signaling in glia is essential for starvation-induced upregulation of Tret1-1.

The Activin-branch of TGF-β signaling has been shown to be important for sugar sensing and sugar metabolism in the adult fly as well as in larvae (Chng et al., 2014; Ghosh and O’Connor, 2014; Mattila et al., 2015). The type I receptor Baboon (Babo) is specific for the Activin branch of TGF-β signaling (reviewed in Upadhyay et al., 2017); Figure 6C). Thus, we silenced *babo* in glial cells using *babo* that has been shown to efficiently abolish *babo* expression (Hevia and de Celis, 2013). Interestingly, in *babo* knockdown animals Tret1-1 expression is strongly upregulated upon starvation (Figure 6), indicating that the Activin-branch of TGF-β signaling is not implicated in Tret1-1 regulation.

This indicates that the BMP-branch of TGF-β signaling is implicated in *tret1-1* regulation. To analyze its involvement, we knocked down the BMP branch-specific type I receptors Thickveins (Tkv) and Saxophone (Sax) (reviewed in Upadhyay et al., 2017). Loss-of-function mutations in both *tkv* and *sax* are lethal, but Tkv overexpression can rescue *sax* loss-of-function, thus Tkv seems to be the primary type I receptor in the BMP-branch of TGF-β signaling (Brummel et al., 1994). Glia-specific knockdown of *sax* using *sax^GD50^* or *sax^GD2546^* did not show any differences in Tret1-1 regulation upon starvation compared to control knockdown animals (Figure 6). In contrast, knockdown of *tkv* using *tkv^KK102319^* abolished Tret1-1 upregulation upon starvation, highlighting its importance for signaling (Figure 6).

### Glass-bottom boat-mediated TGF-β signaling induces Tret1-1 expression upon starvation

The BMP branch of TGF-β signaling can be activated by several ligands, Glass-bottom boat (Gbb), Decapentaplegic (Dpp), Screw (Scw) and probably Maverick (Mav) (reviewed in Upadhyay et al., 2017). Of those ligands only Gbb has been implicated in regulating metabolic processes so far (reviewed in Upadhyay et al., 2017). *gbb^-/-^* mutant animals show a phenotype that resembles the state of starvation, including reduced triacylglyceride storage and lower circulating carbohydrate levels (Ballard et al., 2010). It has previously been shown that overexpression of Gbb in the fatbody leads to higher levels of circulating carbohydrates and thus the opposite of a starvation-like phenotype (Hong et al., 2016a). Thus, to study their role in Tret1-1 regulation, we over-expressed Gbb or Dpp locally in the surface glial cells (9137-Gal4, perineurial and subperineurial glial cells) to avoid strong systemic impact that would counteract the effects of starvation. In fed animals that express Gbb in the BBB-cells Tret1-1 expression is significantly upregulated in the perineurial glial cells (Figure 7). This effect is specific to Gbb, since neither GFP-expressing control animals nor Dpp-expressing animals display this effect (Figure 7). This shows that Gbb-dependent signaling does induce Tret1-1 upregulation.

**Figure 7.**
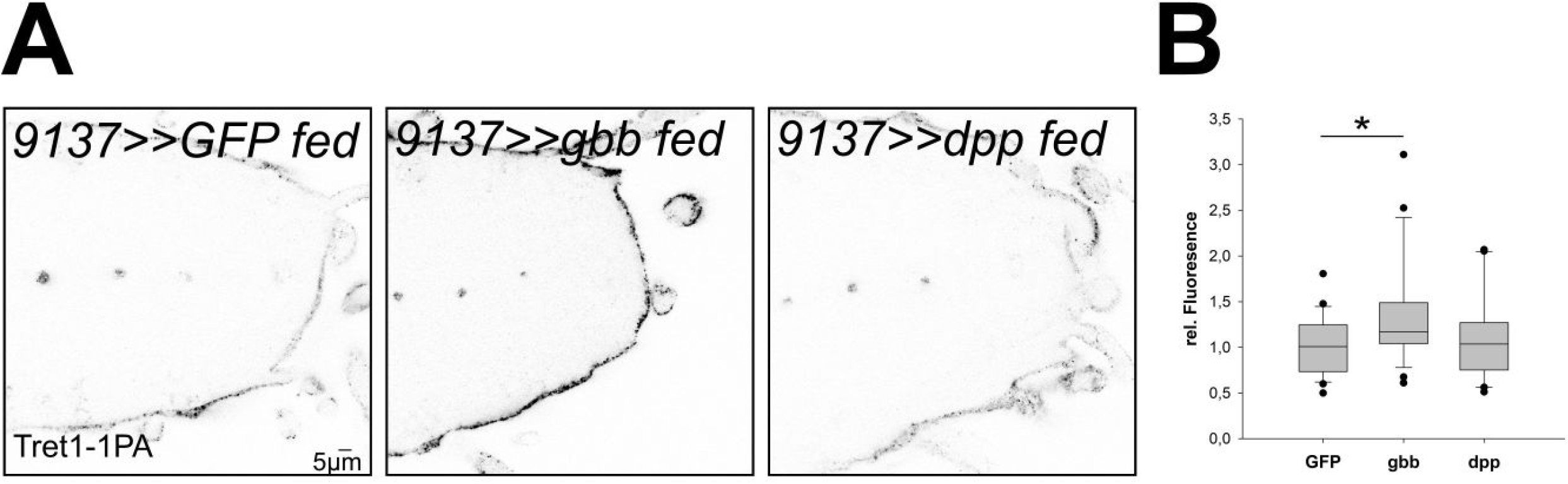

Taken together, the data reported here show that, upon starvation, moderate levels of Gbb are produced by an unknown source, probably locally in the subperineurial glial cells. Gbb activates the BMP-branch of TGF-β signaling in the perineurial glial cells, via the receptors Tkv (type I) and Put (type II) and induces Tret1-1 expression. Since it has been shown that mammalian GLUT1 is also upregulated upon hypoglycemia, it will be interesting to see whether TGF-β signaling is conserved as a pathway adapting carbohydrate transport to changes in nutrient availability.

## Discussion

The nervous system is separated from circulation by the BBB. This separation on one hand protects the nervous system form circulation-derived harmful substances, but on the other hand necessitates efficient nutrient transport to ensure neuronal function. Since the nervous system mainly uses carbohydrates to meet its energetic demands, carbohydrates need to be taken up at a sufficient rate. We previously showed that the carbohydrate transporter Tret1-1 is specifically expressed in perineurial glial cells that surround the Drosophila brain and that glucose is taken up into the nervous system (Volkenhoff et al., 2018, 2015). Here, we investigated how Tret1-1-mediated carbohydrate uptake into the nervous system is adapted to the metabolic state of the animal to spare the nervous system from the effects of malnutrition. We show that Tret1-1 is a carbohydrate transporter that cannot only facilitate transport of trehalose as previously reported (Kanamori et al., 2010), but also of glucose (Figure 4). Upon chronic starvation Tret1-1 protein levels are increased in the perineurial glial cells (Figure 1), boosting the glucose transport capacity in those cells (Figure 4). Even though we cannot exclude additional upregulation of other carbohydrate transporters, the data shown here indicates that Tret1-1 upregulation is a mechanism to ensure efficient carbohydrate uptake, even when circulating carbohydrate levels are low.

Subcellular trafficking of Tret1-1 is important for Tret1-1 homeostasis and its integration into the plasma membrane, which is increased upon starvation (Figure 1, 2). Loss of Rab7 or Rab10 function has severe effects on Tret1-1 levels or localization. The intracellular accumulation of Tret1-1 induced by Rab10 silencing indicates that Tret1-1 cannot be properly delivered to the plasma membrane. Loss of Rab10 function in mammalian adipocytes induces perinuclear accumulation of GLUT4, suggesting regulatory parallels between Tret1-1 and GLUT4 (Sano et al., 2007). GLUT4 (*Slc2a4*) is weakly expressed in the mammalian BBB (James et al., 1988; McCall et al., 1997). Also, the two closest GLUT-homologues of Tret1-1, GLUT6 and GLUT8, are regulated by subcellular trafficking from cytoplasmic storage vesicle to the plasma membrane (FlyBase, Lisinski et al., 2001). Both, GLUT6 and GLUT8 are expressed in the mammalian brain, but their roles are unclear (H. Doege et al., 2000; Holger Doege et al., 2000; Ibberson et al., 2000; Reagan et al., 2002).

We show that the *tret1-1* promoter is induced upon starvation (Figure 3). This suggests that the *tret1-1* locus harbors a starvation-responsive element. Tret1-1 levels are most likely regulated dependent on carbohydrate availability, since animals feeding on sugar-only food do not show an upregulation of Tret1-1 (Figure 1). It has been reported that insulin-induced hypoglycemia leads to an upregulation of GLUT1 mRNA as well as protein in rat BBB-forming endothelial cells (Kumagai et al., 1995). In isolated rat brain microvessels insulin-induced hypoglycemia also activates upregulation of GLUT1 protein levels and in addition an accumulation of GLUT1 at the luminal membrane (Simpson et al., 1999). In these rodent studies, GLUT1 upregulation was detected upon insulin injection that induces hypoglycemia. Under starvation conditions that lead to hypoglycemia in our experimental setup, however, insulin levels are strongly reduced. If under high insulin conditions GLUT1 levels are increased in mammals, this increase cannot be triggered by a loss of insulin. Along the same lines, the upregulation of Tret1-1 in perineurial glial cells we report here is independent of insulin signaling as well as AKH signaling (Figure 5). Thus, the regulatory mechanisms reported here may be conserved. This is especially interesting since aberrations in Glut1 functionality or levels can cause severe diseases, like e.g. Glut1 deficiency syndrome or Alzheimer’s (reviewed in Koepsell, 2020). Therefore, understanding the mechanisms that control the expression of carbohydrate transporters in the BBB-forming cells might be the basis for developing a treatment that allows to correct non-sufficient transporter expression in such diseases.

The induction of carbohydrate transport at the BBB upon hypoglycemia or starvation seems to be a mechanism that is required to spare the brain from the effects of malnutrition. It has previously been shown in mammals, as well as in flies, that the developing nervous system is protected from such effects to allow proper brain growth, while other organs undergo severe growth restriction. This process is called asymmetric intra-uterine growth restriction in humans or “brain sparing” in model organisms (reviewed in Lanet and Maurange, 2014). In Drosophila, the mechanisms that underly the protection of the brain have been studied. Here, Jelly belly (Jeb)/Anaplastic lymphoma kinase (ALK) signaling in the neuroblast niche circumvents the need for insulin signaling to propagate growth (reviewed in Lanet and Maurange, 2014). Interestingly, Jeb/ALK signaling is not the basis for Tret1-1 upregulation in the perineurial glial cells, since glial ALK knockdown does not abolish Tret1-1 induction upon starvation (Figure 6).

TGF-β signaling has been shown to be involved in metabolic regulation in vertebrates and invertebrates (Andersson et al., 2008; Bertolino et al., 2008; Ghosh and O’Connor, 2014; Zamani and Brown, 2011). In Drosophila, the Activin-like ligand Dawdle as well as the BMP ligand Glass-bottom boat have been implicated in metabolic regulation (reviewed in Upadhyay et al., 2017). Daw seems to be one of the primary players in the conserved ChREBP/MondoA-Mlx complex-dependent sugar-sensing pathway (Mattila et al., 2015). However, since the Activin-like branch of TGF-β signaling does not play a role in Tret1-1 regulation, it does not seem to affect carbohydrate uptake into the nervous system. The BMP ligand Gbb, on the other hand, has been implicated in nutrient storage regulation. *gbb* mutants show expression defects of several starvation response genes (Ballard et al., 2010). Furthermore, the fat body of fed *gbb* mutants resembles that of starved wild type animals by its nutrient storage and morphology (Ballard et al., 2010). Gbb seems to be regulating nutrient storage in the fat body and fat body morphology in a cell-autonomous manner, but since *gbb* mutants display increased nutrient uptake rates, *gbb* signaling also has systemic effects that are not yet completely understood (Ballard et al., 2010; Hong et al., 2016b). We show here that moderate levels of Gbb signaling induce an upregulation of Tret1-1 expression in perineurial glial cells (Figure 7). Gbb signals via Tkv and Put to regulate Tret1-1 expression upon starvation (Figure 6). Interestingly, it has been shown that Bmp signaling induces transcriptional upregulation of Glut1 in chondrocytes during murine skeletal development (Lee et al., 2018). Thus, TGF-β dependent regulation of carbohydrate transport at the BBB may be based on the same mechanisms and consequently be evolutionarily conserved.

Interestingly, the transcription of the Drosophila sodium/solute cotransporter cupcake has also been shown to be upregulated upon starvation. Cupcake is expressed in some ellipsoid body neurons upon starvation and is essential for the ability of the animal to choose feeding on a nutritive sugar over feeding on a sweeter non-nutritive sugar after a period of nutrient deprivation. Furthermore, several solute carrier family members have been shown to be regulated by carbohydrate availability in mouse cortical cell culture (Ceder et al., 2020). It will be very interesting to investigate whether such transcriptional upregulation is also mediated by TGF-β signaling and if TGF-β-mediated transcriptional regulation in the nervous system is a central mechanism that allows survival under nutrient shortage.

In summary, we report here a potentially conserved mechanism that spares the nervous system from effects of nutrient shortage by upregulation of carbohydrate transport at the BBB. This upregulation renders carbohydrate uptake more efficient and most likely allows sufficient carbohydrate uptake even when circulating carbohydrate levels are low. In Drosophila, compensatory upregulation of Tret1-1 is regulated via Gbb and the BMP branch of TGF-β signaling. This mechanism is likely to be evolutionarily conserved, since mammalian Glut1 has been shown to be regulated via BMP signaling in other tissues (Lee et al., 2018) and thus might in the future allow designing a treatment against diseases caused by non-sufficient carbohydrate transport in the nervous system.

## Acknowledgement

We are grateful to M. Brankatschk for fly stocks. We thank Astrid Fleige for help with cloning and Western blots. We are grateful to C. Klämbt for discussions and critical reading of the manuscript. The work was supported by grants of the DFG to SS (SFB1009, SCHI 1380/2-1).

## Author contribution

H.H. designed and conducted most experiments, helped conceiving the study and wrote the paper with S.S.; E.M. conducted the FRET experiments and helped writing the paper; A.W. conducted the Xenopus experiments together with H.M.B.; A.V. generated the *tret1-1-Gal4* and *tret1-1-stg-GFP* flies; H.M.B. designed the Xenopus experiments and helped conducting them; S.S. conceived the study, assisted in designing and interpreting experiments, wrote the paper with H.H. and E.M. and obtained funding from the DFG.

## Materials and Methods

All experiments have been conducted at least 3 times independently of each other to assess interexperimental variation. In each experiment several animals have been used to assess variations between animals. N gives the number of independent experiments; n is the total number of animals analyzed. If not noted otherwise, immunostainings have been done 3 times independently including several animals in each experiment.

### Fly stocks

Flies were kept at 25 °C on a standard diet if not noted otherwise. The following fly stocks were used in this study: jeb^KK111857^, jeb^GD5472^, Alk^GD42^, put^KK102676^, put^GD2545^, wit^KK100911^, sax^GD50^, sax^GD2546^, tkv^KK102319^, Rab10^GD13212^, Rab10^GD16778^, Rab10^KK109210^ (all fly stocks were obtained from VDRC fly center). Rab7^T22N^, Rab10^T23N^, Rab7^EYFP^, Rab10^EYFP^, Rab19^EYFP^, Rab23^EYFP^, Rab7^TRIPJF02377^, InR^K1409A^, InR^R418P^, UAS-dpp (BDSC 1486), mCherry^dsRNA^ (BDSC 35785), UAS-CD8-GFP (BDSC 30002 or 30003) (all fly stocks were obtained from Bloomington Drosophila stock center). Akh^AP^ and Akh^SAP^ (Gáliková et al., 2015), babo^NIG8224R^ (Japanese National Institute of Genetics), gliotactin-Gal4, repo-Gal4 (Sepp et al., 2001), 46F-Gal4 (Xie and Auld, 2011), 9137-Gal4 (Desalvo et al., 2014), UAS-FLII^12^Pglu-700μδ6 (Volkenhoff et al., 2018), UAS-Gbb (P. Soba), UAS-RFP (S. Heuser), w^1118^ (Lindsley and Zimm, 1992).

### *Creation of* Tret1-1-Gal4 *and* Tret1-1-stinger-GFP *flies*

For creation of Tret1-1-Gal4 and Tret1-1-stinger-GFP flies, first the promotor region of *tret1-1* was cloned from genomic DNA (forward primer: CACCGGTCTCAAGCTCTCTTTTTTGCCTTACATATTTT, reverse primer: TGGGTAAGTTGGAGAGAGAG) into the pENTR™ vector using the pENTR™/D-TOPO^®^ Cloning Kit (Thermofisher). Via the gateway system, the promotor fragment was cloned either into the pBPGuwGal4 vector (addgene #17575) or into pBPGuw-stingerGFP. Both clones were introduced into the 86Fb landing site via Φ integrase-mediated transgenesis (Bischof et al., 2007).

### Immunohistochemistry, SDS Page and Western blotting

Third instar larval brains or larval brains of animals that had been subjected to the larval starvation protocol, were dissected and immunostained following standard protocols (Volkenhoff et al., 2015). Specimen were analyzed using the Zeiss 710 LSM or the Zeiss 880 LSM and the Airy Scan Module (Zeiss, Oberkochen, Germany). SDS Page and Western blotting was performed following published protocols (Zobel et al., 2015). Lysates were generated from 96 h +/- 3 h old larval brains.

The following antibodies were used: guinea pig anti-Tret1-1 (1:50, Volkenhoff et al., 2015), rabbit anti-Laminin (1:1000, Abcam), mouse anti-Repo (1:2, Developmental Studies Hybridoma Bank), mouse anti-GFP (for immunohistochemistry 1:1000, Molecular Probes; for Western blotting: 1:10000, Clontech), mouse anti-Tubulin (1:80, Developmental Studies Hybridoma Bank), rabbit anti-Apontic (1:150, Eulenberg and Schuh, 1997). As secondary conjugated antibodies, Alexa488- (1:1000), Alexa568- (1:1000) and Alexa647-coupled (1:500) antibodies were used (all from Thermo Fisher Scientific). For Western blotting, goat anti-mouse HRP (Dianova, 1:7500) was used. HRP activity was detected using the ECL detection system kit (GE Healthcare) and the Amersham Imager 680 (GE Healthcare). Image analysis was performed using the Fiji plugin of ImageJ (1.52p, java 1.8.0._172 64-bit, NIH, Bethesda, Maryland).

### Larval starvation

Flies were kept overnight on standard food to stage the embryos. 42 h after staging similar sized larvae were collected, cleared from food and transferred to different food conditions: standard food, water-soaked filter paper or 10 % sucrose in PBS. They were kept for 48 h on this condition before dissecting. For fluorescent analysis, mean grey values of a region of interest (ROI) containing the entire tip of the ventral nerve cord were measured. The mean of values of seven single planes was taken. To obtain comparable values between experiments, the ratio of values received from starved animals to fed animals was calculated. Statistical analysis was performed using Sigma Plot software (Jadel). Differences were assessed by the Mann-Whitney Rank Sum test or t-test. P values < 0.05 were considered as significantly different.

### Measurement of glucose uptake

Larvae expressing *UAS-FLII^12^Pglu-700μδ6* FRET glucose sensor under the control of 9137-Gal4 were kept on standard food or under starvation conditions following the larval starvation protocol. Larval brains were subsequently dissected in HL3 buffer (70 mM NaCl, 5 mM KCl, 20 mM MgCl_2_, 10 mM NaHCO_3_, 115 mM sucrose, 5mM trehalose, 5 mM HEPES; pH 7.2; ca. 350 mOsm) and adhered to Poly-D-Lysine-coated coverslips. Coverslips were secured into a flow through chamber and mounted to the stage of a LSM880 confocal microscope (Zeiss, Oberkochen, Germany). The chamber was then connected to a mini-peristaltic pump (MPII, Harvard Apparatus) to allow buffer exchange.

Fluorescent images were acquired immediately after dissection using 20x/1,0 DIC M27 75mm emersion objective (Zeiss, Oberkochen, Germany) with excitation 436/25 nm, beam splitter 455 nm, emission 480/40 nm (CFP channel); excitation 436/25 nm, beam splitter 455 nm, emission 535/30 nm (YFP channel). Each larval brain was imaged in a separate experiment (n=10). After 2.5 minutes, HL3 buffer was exchanged for glucose buffer (HL3 supplemented with 10 mM glucose; pH 7.2) and replaced by HL3 again after a further 7.5 minutes.

For data analysis, a ROI containing the entire larval brain was selected and the mean grey value of all pixels minus background for each channel was calculated. Values were normalized to known minimum (HL3 buffer). Statistical and regression analysis of data obtained was performed using SigmaPlot software (Jandel). To determine glucose uptake rates, 10 time points 9 seconds after values rose above baseline levels were used to calculate the linear slope of each curve. Differences were assessed by the Mann-Whitney Rank Sum test (pairs). P values < 0.05 were considered as significantly different.

### Xenopus experiments

For isolation of oocytes, female *X. laevis* frogs (purchased from the Radboud University, Nijmegen, Netherlands) were anesthetized with 1 g/l of Ethyl 3-aminobenzoate methanesulfonate and rendered hypothermic. Parts of ovarian lobules were surgically removed under sterile conditions. The procedure was approved by the Landesuntersuchungsamt Rheinland-Pfalz, Koblenz (23 177-07/A07-2-003 §6). Oocytes were singularized by collagenase treatment in Ca^2+^-free oocyte saline (82.5 mM NaCl, 2.5 mM KCl, 1 mM MgCl_2_, 1 mM Na_2_HPO_4_, 5 mM HEPES, pH 7.8, 2 mg/l gentamicin) at 28 °C for 2 h. The singularized oocytes were stored overnight at 18 °C in Ca^2+^-containing oocyte saline (82.5 mM NaCl, 2.5 mM KCl, 1 mM CaCl2, 1 mM MgCl2, 1 mM Na_2_HPO_4_, 5 mM HEPES, pH 7.8, 2 mg/l gentamicin). The procedure was described in detail previously (Becker, 2014).

For heterologous protein expression in *X. laevis* oocytes the *D. melanogaster* cDNA sequences of Tret1-1 isoform A was amplified via PCR from pUAST-Tret1-1-PA-3xHA plasmid (forward primer: CGTCTAGAATGAGTGGACGCGAC, reverse primer: CGAAGCTTCTAGCTTACGTCACGT) and cloned into the pGEM-He-Juel vector using XbaI/ HindIII restriction sites. cRNA was produced by *in vitro* transcription using the mMESSAGE mMACHINE^®^ T7 Kit (Fisher Scientific). Oocytes of the stages V and VI were injected with 18 ng (for mass spectrometry) to 20 ng (for scintillation analysis) of cRNA and measurements were carried out three to six days after cRNA injection.

To analyze the transport capacity by scintillation measurements radioactive sugar substrates were generated using unlabeled sugar solutions of different concentrations in oocyte saline and adding ^14^C-labeled sugar at a concentration of 0.15 μCi/100 μl (for 0.3 mM to 30 mM solutions) or 0.3 μCi/100 μl (for 100 mM and 300 mM solutions). ^14^C_12_-trehalose was purchased from Hartmann Analytic, Braunschweig (#1249), ^14^C_6_-glucose and ^14^C_6_-fructose were purchased from Biotrend, Köln (#MC144-50 and 66 #MC1459-50). Six to eight oocytes were transferred into a test tube and washed with oocyte saline. Oocyte saline was removed completely and 95 μl of the sugar substrate were added for 60 min. After incubation, cells were washed four times with 4 ml ice-cold oocyte saline. Single oocytes were transferred into Pico Prias scintillation vials (Perkin Elmer) and lysed in 200 μl 5 % SDS, shaking at approximately 190 rpm for at least 30 min at 20 to 28 °C. 3 ml Rotiszint^®^ eco plus scintillation cocktail (Carl Roth) were added to each vial and scintillation was measured using the Tri-Carb 2810TR scintillation counter (Perkin Elmer). Scintillation of 10 μl sugar substrate of each concentration with 200 μl 5 % SDS and 3 ml Rotiszint^®^ eco plus scintillation cocktail served as a standard.

Substrate flux was calculated from the measured scintillation according to the respective standard measurements. For statistical analysis, the medium flux and standard error were calculated for oocytes expressing transport proteins and native oocytes and compared using a one-sided t-test or the Man-Whitney Rank test for analysis of non-uniformly distributed samples. Determination of the net-flux was performed by subtracting the medium flux of native oocytes from one test series from each measurement of the same test series and calculating the medium flux and standard error.

## Supplementary information

**Figure S1.**
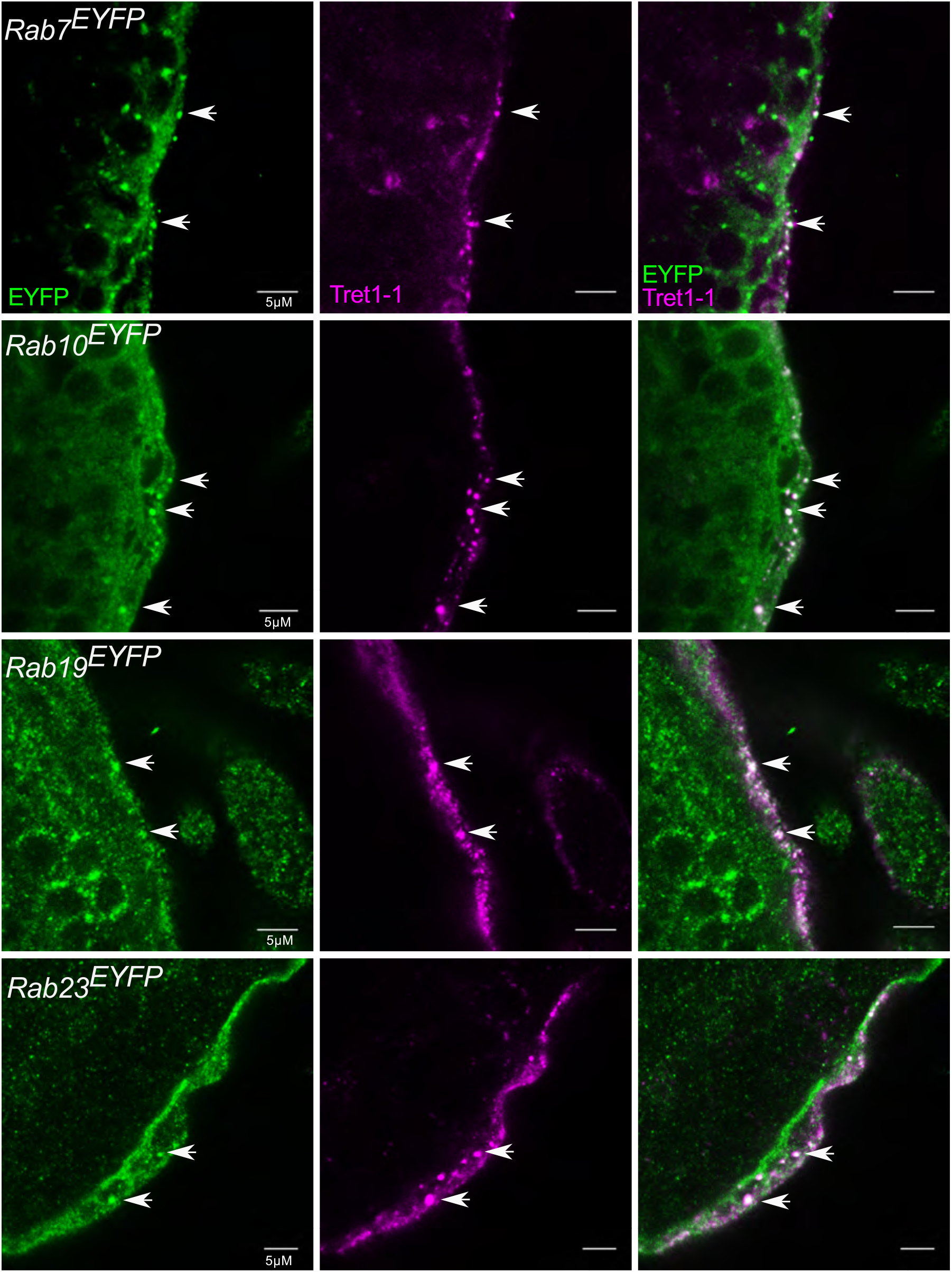
Rab7, Rab10, Rab19 and Rab23 colocalize with Tret1-1 vesicles. Co-staining of endogenous EYFP-tagged Rab-GTPases (green) and Tret1-1 (magenta) in the surface glia of third instar larval brains. All Drosophila Rab-GTPases endogenously labeled with EYFP were tested. Tret1-1-positive vesicles show overlapping staining with Rab7^EYFP^, Rab10^EYFP^, Rab19^EYFP^ and Rab23^EYFP^.

**Figure S2.**
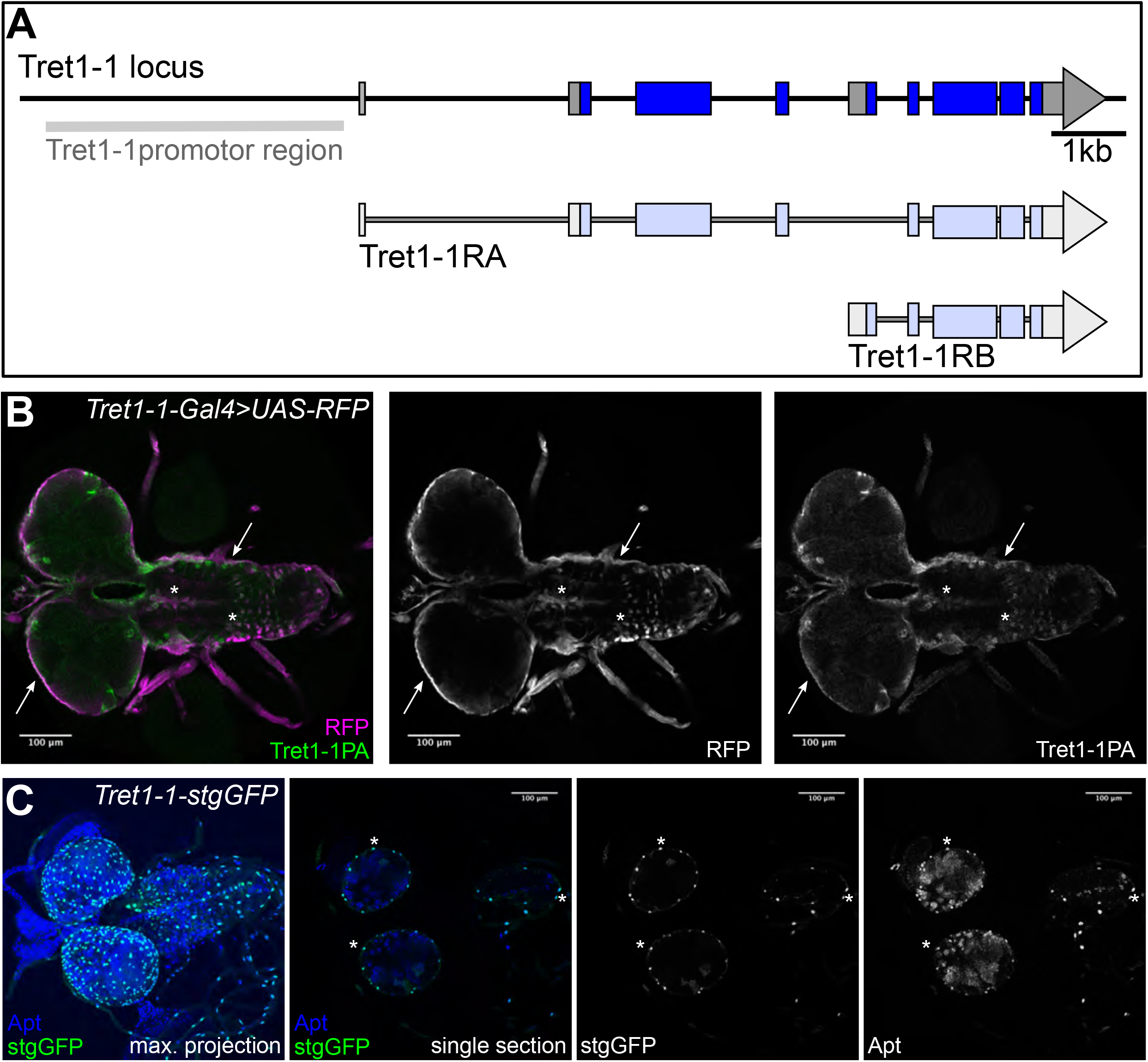
Tret1-1 promoter drives specific expression. **(A)** Schematic of the *tret1-1* locus and the transcripts encoding the two Tret1-1 isoforms. The *tret1-1* promoter region used to generate *tret1-1-Gal4* and *tret1-1-stgGFP* is highlighted in grey. **(B)** Tret1-1PA staining (green, grey) overlaps with *tret1-1* -driven RFP (magenta, grey) *(tret1-1-Gal4 UAS-RFP)*, verifying the specificity of the *tret1-1* promotor region. **(C)** Co-staining of Apontic (blue, grey) and stgGFP (green, grey) that shows that *tret1-1*-driven stgGFP is specifically expressed in perineurial glial nuclei.

**Figure S3.**
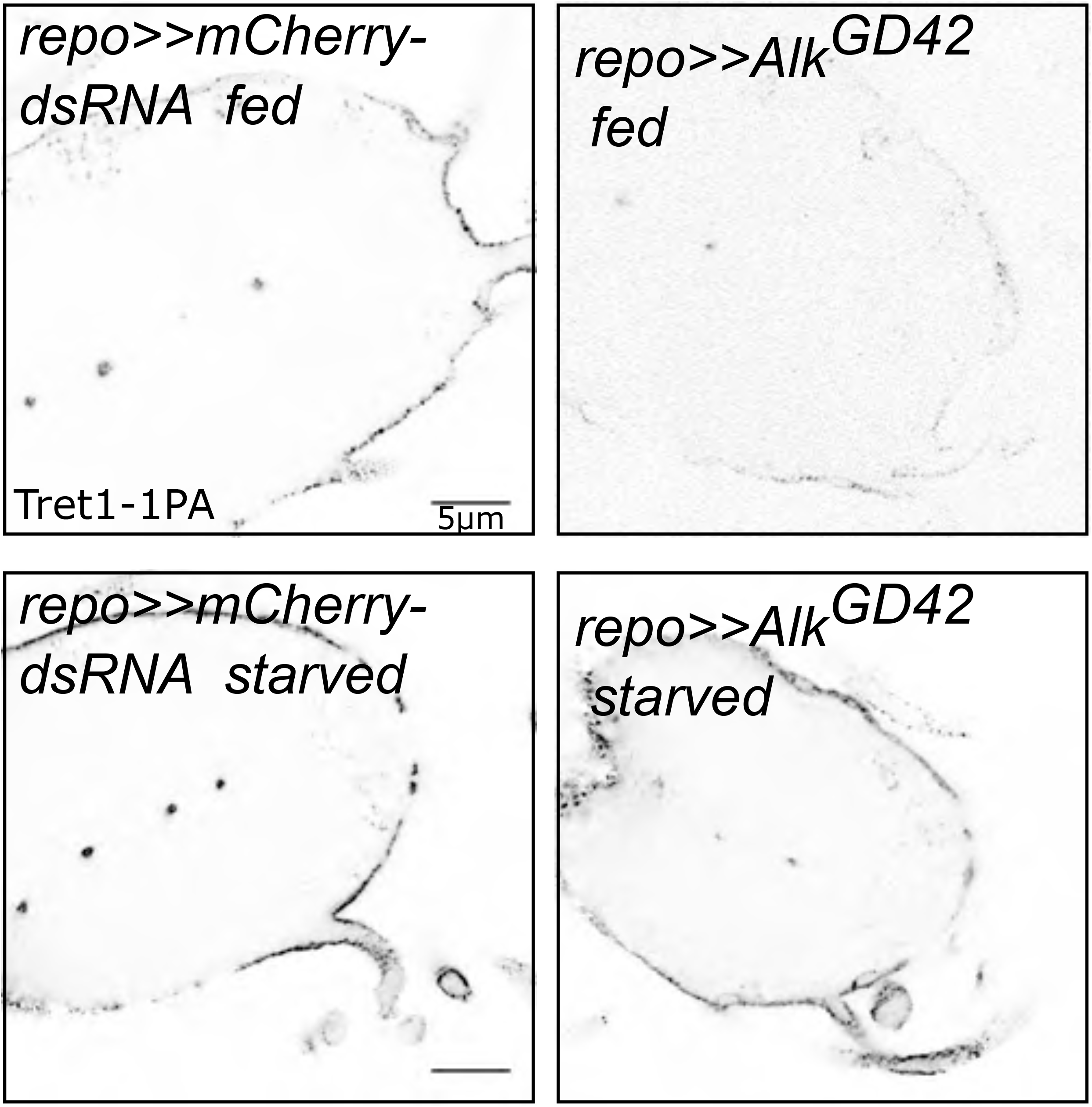
Tret1-1 regulation upon starvation is ALK-independent. Tret1-1 staining of the ventral nerve cord of starved and fed control (*repo>>mCherry-dsRNA*) and *alk* knockdown (*repo>>alk^GD42^*) animals. Tret1-1 upregulation is still induced in starved *alk* knockdown animals.

## References

Andersson O, Korach-Andre M, Reissmann E, Ibáñez CF, Bertolino P. 2008. Growth/differentiation factor 3 signals through ALK7 and regulates accumulation of adipose tissue and diet-induced obesity. Proc Natl Acad Sci U S A 105:7252–7256. doi:10.1073/pnas.0800272105

Arsov T, Mullen SA, Damiano JA, Lawrence KM, Huh LL, Nolan M, Young H, Thouin A, Dahl H-HM, Berkovic SF, Crompton DE, Sadleir LG, Scheffer IE. 2012. Early onset absence epilepsy: 1 in 10 cases is caused by GLUT1 deficiency. Epilepsia 53:e204–e207. doi:10.1111/epi.12007

Ballard SL, Jarolimova J, Wharton KA. 2010. Gbb/BMP signaling is required to maintain energy homeostasis in Drosophila. Dev Biol 337:375–385.

Begley DJ. 2006. Structure and function of the blood-brain barrierEnhancement in Drug Delivery. doi:10.3389/conf.fphar.2010.02.00002

Bertolino P, Holmberg R, Reissmann E, Andersson O, Berggren PO, Ibáñez CF. 2008. Activin B receptor ALK7 is a negative regulator of pancreatic β-cell function. Proc Natl Acad Sci U S A 105:7246–7251. doi:10.1073/pnas.0801285105

Blatt J, Roces F. 2001. Haemolymph sugar levels in foraging honeybees (Apis mellifera carnica): dependence on metabolic rate and in vivo measurement of maximal rates of trehalose synthesis. J Exp Biol 204:2709–16.

Boado RJ, Pardridge WM. 1993. Glucose Deprivation Causes Posttranscriptional Enhancement of Brain Capillary Endothelial Glucose Transporter Gene Expression via GLUT1 mRNA Stabilization. J Neurochem 60:2290–2296. doi: 10.1111/j.1471-4159.1993.tb03516.x

Broughton S, Alic N, Slack C, Bass T, Ikeya T, Vinti G, Tommasi AM, Driege Y, Hafen E, Partridge L. 2008. Reduction of DILP2 in Drosophila triages a metabolic phenotype from lifespan revealing redundancy and compensation among DILPs. PLoS One 3:e3721.

Brummel TJ, Twombly V, Marqués G, Wrana JL, Newfeld SJ, Attisano L, Massagué J, O’Connor MB, Gelbart WM. 1994. Characterization and relationship of dpp receptors encoded by the saxophone and thick veins genes in Drosophila. Cell 78:251–261. doi:10.1016/0092-8674(94)90295-X

Ceder MM, Lekholm E, Klaesson A, Tripathi R, Schweizer N, Weldai L, Patil S, Fredriksson R. 2020. Glucose Availability Alters Gene and Protein Expression of Several Newly Classified and Putative Solute Carriers in Mice Cortex Cell Culture and D. melanogaster. Front Cell Dev Biol 8:579. doi:10.3389/fcell.2020.00579

Cheng LY, Bailey AP, Leevers SJ, Ragan TJ, Driscoll PC, Gould AP. 2011. Anaplastic Lymphoma Kinase Spares Organ Growth during Nutrient Restriction in Drosophila. Cell 146:435–447. doi:10.1016/j.cell.2011.06.040

Chng WA, Sleiman MSB, Schüpfer F, Lemaitre B. 2014. Transforming Growth Factor β/Activin Signaling Functions as a Sugar-Sensing Feedback Loop to Regulate Digestive Enzyme Expression. Cell Rep 9:336–348. doi:https://doi.org/10.1016/j.celrep.2014.08.064

Çiçek IÖ, Karaca S, Brankatschk M, Eaton S, Urlaub H, Shcherbata HR. 2016. Hedgehog Signaling Strength Is Orchestrated by the *mir-310* Cluster of MicroRNAs in Response to Diet. Genetics 202:1167 LP – 1183. doi:10.1534/genetics.115.185371

Corvera S, Chawla A, Chakrabarti R, Joly M, Buxton J, Czech MP. 1994. A double leucine within the GLUT4 glucose transporter COOH-terminal domain functions as an endocytosis signal. J Cell Biol 126:979–989. doi:10.1083/jcb.126.4.979

Croset V, Treiber CD, Waddell S. 2018. Cellular diversity in the Drosophila midbrain revealed by single-cell transcriptomics. Elife 7:e34550. doi:10.7554/eLife.34550

Cushman SW, Wardzala LJ. 1980. Potential mechanism of insulin action on glucose transport in the isolated rat adipose cell. Apparent translocation of intracellular transport systems to the plasma membrane. J Biol Chem 255:4758–4762.

Davie K, Janssens J, Koldere D, De Waegeneer M, Pech U, Kreft Ł, Aibar S, Makhzami S, Christiaens V, Bravo González-Blas C, Poovathingal S, Hulselmans G, Spanier KI, Moerman T, Vanspauwen B, Geurs S, Voet T, Lammertyn J, Thienpont B, Liu S, Konstantinides N, Fiers M, Verstreken P, Aerts S. 2018. A Single-Cell Transcriptome Atlas of the Aging Drosophila Brain. Cell 174:982–998.e20. doi:10.1016/j.cell.2018.05.057

Desalvo MK, Hindle SJ, Rusan ZM, Orng S, Eddison M, Halliwill K, Bainton RJ. 2014. The Drosophila surface glia transcriptome: evolutionary conserved blood-brain barrier processes. Front Neurosci 8:346.

Doege H., Bocianski A, Joost HG, Schurmann A. 2000. Activity and genomic organization of human glucose transporter 9 (GLUT9), a novel member of the family of sugar-transport facilitators predominantly expressed in brain and leucocytes. Biochem J 350 Pt 3:771–776. doi:10.1042/0264-6021:3500771

Doege Holger, Schürmann A, Bahrenberg G, Brauers A, Joost HG. 2000. GLUT8, a novel member of the sugar transport facilitator family with glucose transport activity. J Biol Chem 275:16275–16280. doi:10.1074/jbc.275.21.16275

Dunst S, Kazimiers T, von Zadow F, Jambor H, Sagner A, Brankatschk B, Mahmoud A, Spannl S, Tomancak P, Eaton S, Brankatschk M. 2015. Endogenously tagged rab proteins: a resource to study membrane trafficking in Drosophila. Dev Cell 33:351–365.

Dus M, Min S, Keene AC, Lee GY, Suh GSB. 2011. Taste-independent detection of the caloric content of sugar in <em>Drosophila</em>. Proc Natl Acad Sci 108:11644 LP – 11649.

Elfeber K, Köhler A, Lutzenburg M, Osswald C, Galla HJ, Witte OW, Koepsell H. 2004. Localization of the Na+-D-glucose cotransporter SGLT1 in the blood-brain barrier. Histochem Cell Biol 121:201–207. doi:10.1007/s00418-004-0633-9

Enerson BE, Drewes LR. 2006. The rat blood-brain barrier transcriptome. J Cereb Blood Flow Metab 26:959–973. doi:10.1038/sj.jcbfm.9600249

Eulenberg KG, Schuh R. 1997. The tracheae defective gene encodes a bZIP protein that controls tracheal cell movement during Drosophila embryogenesis. EMBO J 16:7156–7165. doi:10.1093/emboj/16.23.7156

Gáliková M, Diesner M, Klepsatel P, Hehlert P, Xu Y, Bickmeyer I, Predel R, Kühnlein RP. 2015. Energy Homeostasis Control in Drosophila Adipokinetic Hormone Mutants. Genetics 201:665–683. doi:10.1534/genetics.115.178897

Ghosh AC, O’Connor MB. 2014. Systemic Activin signaling independently regulates sugar homeostasis, cellular metabolism, and pH balance in Drosophila melanogaster. Proc Natl Acad Sci U S A 111:5729–5734. doi:10.1073/pnas.1319116111

Guerra F, Bucci C. 2016. Multiple Roles of the Small GTPase Rab7. Cells 5:34. doi:10.3390/cells5030034

Harris JJ, Jolivet R, Attwell D. 2012. Synaptic energy use and supply. Neuron 75:762–777.

Hevia CF, de Celis JF. 2013. Activation and function of TGFβ signalling during Drosophila wing development and its interactions with the BMP pathway. Dev Biol 377:138–153. doi:https://doi.org/10.1016/j.ydbio.2013.02.004

Hindle SJ, Bainton RJ. 2014. Barrier mechanisms in the Drosophila blood-brain barrier. Front Neurosci 8:414.

Ho T-Y, Wu W-H, Hung S-J, Liu T, Lee Y-M, Liu Y-H. 2019. Expressional Profiling of Carpet Glia in the Developing Drosophila Eye Reveals Its Molecular Signature of Morphology Regulators. Front Neurosci 13:244. doi:10.3389/fnins.2019.00244

Hoffmann U, Sukhotinsky I, Eikermann-Haerter K, Ayata C. 2013. Glucose modulation of spreading depression susceptibility. J Cereb Blood Flow Metab 33:191–195. doi:10.1038/jcbfm.2012.132

Hong S-H, Kang M, Lee K-S, Yu K. 2016a. High fat diet-induced TGF-β/Gbb signaling provokes insulin resistance through the tribbles expression. Sci Rep 6:30265. doi:10.1038/srep30265

Hong S-H, Kang M, Lee K-S, Yu K. 2016b. High fat diet-induced TGF-β/Gbb signaling provokes insulin resistance through the tribbles expression. Sci Rep 6:30265. doi:10.1038/srep30265

Huotari J, Helenius A. 2011. Endosome maturation. EMBO J 30:3481–3500. doi:10.1038/emboj.2011.286

Ibberson M, Uldry M, Thorens B. 2000. GLUTX1, a novel mammalian glucose transporter expressed in the central nervous system and insulin-sensitive tissues. J Biol Chem 275:4607–4612. doi:10.1074/jbc.275.7.4607

James DE, Brown R, Navarro J, Pilch PF. 1988. Insulin-regulatable tissues express a unique insulin-sensitive glucose transport protein. Nature 333:183–185. doi:10.1038/333183a0

Kanamori Y, Saito A, Hagiwara-Komoda Y, Tanaka D, Mitsumasu K, Kikuta S, Watanabe M, Cornette R, Kikawada T, Okuda T. 2010. The trehalose transporter 1 gene sequence is conserved in insects and encodes proteins with different kinetic properties involved in trehalose import into peripheral tissues. Insect Biochem Mol Biol 40:30–37.

Kapogiannis D, Mattson MP. 2011. Disrupted energy metabolism and neuronal circuit dysfunction in cognitive impairment and Alzheimer’s disease. Lancet Neurol 10:187–198. doi:10.1016/S1474-4422(10)70277-5

Klip A, McGraw TE, James DE. 2019. Thirty sweet years of GLUT4. J Biol Chem 294:11369–11381. doi:10.1074/jbc.REV119.008351

Koepsell H. 2020. Glucose transporters in brain in health and disease. Pflügers Arch - Eur J Physiol. doi:10.1007/s00424-020-02441-x

Kumagai AK, Kang Y-S, Boado RJ, Pardridge WM. 1995. Upregulation of Blood-Brain Barrier GLUT1 Glucose Transporter Protein and mRNA in Experimental Chronic Hypoglycemia. Diabetes 44:1399 LP – 1404. doi:10.2337/diab.44.12.1399

Kuzawa CW, Chugani HT, Grossman LI, Lipovich L, Muzik O, Hof PR, Wildman DE, Sherwood CC, Leonard WR, Lange N. 2014. Metabolic costs and evolutionary implications of human brain development. Proc Natl Acad Sci U S A 111:13010–13015. doi:10.1073/pnas.1323099111

Lane NJ, Treherne JE. 1972. Studies on perineurial junctional complexes and the sites of uptake of microperoxidase and lanthanum in the cockroach central nervous system. Tissue Cell 4:427–436. doi:https://doi.org/10.1016/S0040-8166(72)80019-3

Lanet E, Maurange C. 2014. Building a brain under nutritional restriction: insights on sparing and plasticity from Drosophila studies. Front Physiol 5:117. doi:10.3389/fphys.2014.00117

Laughlin SB, de Ruyter Van Steveninck RR, Anderson JC. 1998. The metabolic cost of neural information. Nat Neurosci 1:36–41. doi:10.1038/236

Lee G, Park JH. 2004. Hemolymph Sugar Homeostasis and Starvation-Induced Hyperactivity Affected by Genetic Manipulations of the Adipokinetic Hormone-Encoding Gene in Drosophila melanogaster. Genetics 167:311–323.

Lee S-Y, Abel ED, Long F. 2018. Glucose metabolism induced by Bmp signaling is essential for murine skeletal development. Nat Commun 9:4831. doi:10.1038/s41467-018-07316-5

Limmer S, Weiler A, Volkenhoff A, Babatz F, Klämbt C. 2014. The Drosophila blood-brain barrier: development and function of a glial endothelium. Front Neurosci 8:365.

Lindsley DL, Zimm GG. 1992. The Genome of Drosophila Melanogaster. Elsevier Science.

Lisinski I, Schürmann A, Joost HG, Cushman SW, Al-Hasani H. 2001. Targeting of GLUT6 (formerly GLUT9) and GLUT8 in rat adipose cells. Biochem J 358:517–522. doi:10.1042/0264-6021:3580517

Matsuda H, Yamada T, Yoshida M, Nishimura T. 2015. Flies without trehalose. J Biol Chem 290:1244–1255. doi:10.1074/jbc.M114.619411

Mattila J, Havula E, Suominen E, Teesalu M, Surakka I, Hynynen R, Kilpinen H, Väänänen J, Hovatta I, Käkelä R, Ripatti S, Sandmann T, Hietakangas V. 2015. Mondo-Mlx Mediates Organismal Sugar Sensing through the Gli-Similar Transcription Factor Sugarbabe. Cell Rep 132:350–364. doi:10.1016/j.celrep.2015.08.081

Mayer CM, Belsham DD. 2009. Insulin directly regulates NPY and AgRP gene expression via the MAPK MEK/ERK signal transduction pathway in mHypoE-46 hypothalamic neurons. Mol Cell Endocrinol 307:99–108.

McCall AL, Van Bueren AM, Huang L, Stenbit A, Celnik E, Charron MJ. 1997. Forebrain endothelium expresses GLUT4, the insulin-responsive glucose transporter. Brain Res 744:318–326. doi:10.1016/S0006-8993(96)01122-5

Mink JW, Blumenschine RJ, Adams DB. 1981. Ratio of central nervous system to body metabolism in vertebrates: Its constancy and functional basis. Am J Physiol - Regul Integr Comp Physiol 241:R203–R212. doi:10.1152/ajpregu.1981.241.3.r203

Nagy P, Szatmári Z, Sándor GO, Lippai M, Hegedűs K, Juhász G. 2017. *Drosophila* Atg16 promotes enteroendocrine cell differentiation via regulation of intestinal Slit/Robo signaling. Development 144:3990 LP – 4001. doi:10.1242/dev.147033

Nässel DR, Liu Y, Luo J. 2015. Insulin/IGF signaling and its regulation in Drosophila. Gen Comp Endocrinol 221:255–266. doi:https://doi.org/10.1016/j.ygcen.2014.11.021

Nishizaki T, Kammesheidt A, Sumikawa K, Asada T, Okada Y. 1995. A sodium- and energy-dependent glucose transporter with similarities to SGLT1-2 is expressed in bovine cortical vessels. Neurosci Res 22:13–22. doi:10.1016/0168-0102(95)00876-U

Nishizaki T, Matsuoka T. 1998. Low glucose enhances Na+/glucose transport in bovine brian artery endothelial cells. Stroke 29:844–849. doi:10.1161/01.STR.29.4.844

Pasco MY, Léopold P. 2012. High sugar-induced insulin resistance in Drosophila relies on the lipocalin Neural Lazarillo. PLoS One 7:e36583.

Pataki C, Matusek T, Kurucz É, Andó I, Jenny A, Mihály J. 2010. Drosophila Rab23 Is Involved in the Regulation of the Number and Planar Polarization of the Adult Cuticular Hairs. Genetics 184:1051 LP – 1065. doi:10.1534/genetics.109.112060

Patching SG. 2016. Glucose Transporters at the Blood-Brain Barrier: Function, Regulation and Gateways for Drug Delivery. Mol Neurobiol 54:1046–1077.

Reagan LP, Rosell DR, Alves SE, Hoskin EK, McCall AL, Charron MJ, McEwen BS. 2002. GLUT8 glucose transporter is localized to excitatory and inhibitory neurons in the rat hippocampus. Brain Res 932:129–134. doi:10.1016/S0006-8993(02)02308-9

Rehni AK, Dave KR. 2018. Impact of Hypoglycemia on Brain Metabolism During Diabetes. Mol Neurobiol. doi:10.1007/s12035-018-1044-6

Sano H, Eguez L, Teruel MN, Fukuda M, Chuang TD, Chavez JA, Lienhard GE, McGraw TE. 2007. Rab10, a Target of the AS160 Rab GAP, Is Required for Insulin-Stimulated Translocation of GLUT4 to the Adipocyte Plasma Membrane. Cell Metab 5:293–303. doi:10.1016/j.cmet.2007.03.001

Simpson IA, Appel NM, Hokari M, Oki J, Holman GD, Maher F, Koehler-Stec EM, Vannucci SJ, Smith QR. 1999. Blood-brain barrier glucose transporter: Effects of hypo- and hyperglycemia revisited. J Neurochem. doi:10.1046/j.1471-4159.1999.0720238.x

Stork T, Engelen D, Krudewig A, Silies M, Bainton RJ, Klambt C. 2008. Organization and Function of the Blood Brain Barrier in Drosophila. J Neurosci 28:587–597.

Suzuki K, Kono T. 1980. Evidence that insulin causes translocation of glucose transport activity to the plasma membrane from an intracellular storage site. Proc Natl Acad Sci U S A 77:2542–2545. doi:10.1073/pnas.77.5.2542

Takanaga H, Chaudhuri B, Frommer WB. 2008. GLUT1 and GLUT9 as major contributors to glucose influx in HepG2 cells identified by a high sensitivity intramolecular FRET glucose sensor. Biochim Biophys Acta - Biomembr 1778:1091–1099. doi:10.1016/j.bbamem.2007.11.015

Upadhyay A, Moss-Taylor L, Kim M-J, Ghosh AC, O’Connor MB. 2017. TGF-β Family Signaling in Drosophila. Cold Spring Harb Perspect Biol 9.

van de Waterbeemd H, Camenisch G, Folkers G, Chretien JR, Raevsky OA. 1998. Estimation of blood-brain barrier crossing of drugs using molecular size and shape, and H-bonding descriptors. J Drug Target 151–165. doi:10.3109/10611869808997889

Vemula S, Roder KE, Yang T, Bhat GJ, Thekkumkara TJ, Abbruscato TJ. 2009. A functional role for sodium-dependent glucose transport across the blood-brain barrier during oxygen glucose deprivation. J Pharmacol Exp Ther 328:487–95. doi:10.1124/jpet.108.146589

Volkenhoff A, Hirrlinger J, Kappel JM, Klämbt C, Schirmeier S. 2018. Live imaging using a FRET glucose sensor reveals glucose delivery to all cell types in the Drosophila brain. J Insect Physiol 106:55–64. doi:https://doi.org/10.1016/j.jinsphys.2017.07.010

Volkenhoff A, Weiler A, Letzel M, Stehling M, Klämbt C, Schirmeier S. 2015. Glial Glycolysis Is Essential for Neuronal Survival in Drosophila. Cell Metab 22:437–447. doi:10.1016/j.cmet.2015.07.006

Weiler A, Volkenhoff A, Hertenstein H, Schirmeier S. 2017. Metabolite transport across the mammalian and insect brain diffusion barriers. Neurobiol Dis. doi:10.1016/j.nbd.2017.02.008

Weiszmann R, Hammonds AS, Celniker SE. 2009. Determination of gene expression patterns using high-throughput RNA in situ hybridization to whole-mount Drosophila embryos. Nat Protoc 4:605–618.

Wyatt GR, Kalf GF. 1957. The chemistry of insect hemolymph: II. Trehalose and other carbohydrates. J Gen Physiol 40:833–847.

Yildirim K, Petri J, Kottmeier R, Klämbt C. 2019. Drosophila glia: Few cell types and many conserved functions. Glia 67:5–26. doi:10.1002/glia.23459

Yu AS, Hirayama BA, Timbol G, Liu J, Diez-Sampedro A, Kepe V, Satyamurthy N, Huang SC, Wright EM, Barrio JR. 2013. Regional distribution of SGLT activity in rat brain in vivo. Am J Physiol - Cell Physiol 304:C240–C247. doi:10.1152/ajpcell.00317.2012

Zamani N, Brown CW. 2011. Emerging roles for the transforming growth factor-β superfamily in regulating adiposity and energy expenditure. Endocr Rev. doi:10.1210/er.2010-0018

Zinke I, Schütz CS, Katzenberger JD, Bauer M, Pankratz MJ. 2002. Nutrient control of gene expression in Drosophila: microarray analysis of starvation and sugar-dependent response. EMBO J 21:6162–6173.

Zobel T, Brinkmann K, Koch N, Schneider K, Seemann E, Fleige A, Qualmann B, Kessels MM, Bogdan S. 2015. Cooperative functions of the two F-BAR proteins Cip4 and Nostrin in the regulation of E-cadherin in epithelial morphogenesis. J Cell Sci 128:499 LP – 515. doi:10.1242/jcs.155929

Zülbahar S, Sieglitz F, Kottmeier R, Altenhein B, Rumpf S, Klämbt C. 2018. Differential expression of öbek controls ploidy in the Drosophila blood-brain barrier. Dev 145. doi:10.1242/dev.164111

